# *foxr1* is a novel maternal-effect gene in fish that regulates embryonic cell growth via *p21* and *rictor*

**DOI:** 10.1101/294785

**Authors:** Caroline T. Cheung, Amélie Patinote, Yann Guiguen, Julien Bobe

## Abstract

The family of forkhead box (Fox) transcription factors regulate gonadogenesis and embryogenesis, but the role of *foxr1*/*foxn5* in reproduction is unknown. Evolution of *foxr1* in vertebrates was examined and the gene found to exist in most vertebrates, including mammals, ray-finned fish, amphibians, and sauropsids. By quantitative PCR and RNA-seq, we found that *foxr1* had an ovarian-specific expression in zebrafish, a common feature of maternal-effect genes. In addition, it was demonstrated using *in situ* hybridization that *foxr1* was a maternally-inherited transcript that was highly expressed even in early-stage oocytes and accumulated in the developing eggs during oogenesis. We also analyzed the function of *foxr1* in female reproduction using a zebrafish CRISPR/Cas9 knockout model. It was observed that embryos from the *foxr1*-deficient females had a significantly lower survival rate whereby they either failed to undergo cell division or underwent abnormal division that culminated in growth arrest at around the mid-blastula transition and early death. These mutant-derived eggs contained a dramatically increased level of *p21*, a cell cycle inhibitor, and reduced *rictor*, a component of mTOR and regulator of cell survival, which were in line with the observed growth arrest phenotype. Our study shows for the first time that *foxr1* is an essential maternal-effect gene and is required for proper cell division and survival via the p21 and mTOR pathways. These novel findings will broaden our knowledge on the functions of specific maternal factors stored in the developing egg and the underlying mechanisms that contribute to reproductive fitness.

**Summary sentence:** The *foxr1* gene in zebrafish is a novel maternal-effect gene that is required for proper cell division in the earliest stage of embryonic development possibly as a transcriptional factor for cell cycle progression regulators, *p21* and *rictor*.

## Introduction

In vertebrates, maternal products including transcripts, proteins, and other biomolecules are necessary for kick-starting early embryonic development until the mid-blastula transition (MBT) when the zygotic genome is activated [1]. Maternal-effect genes are transcribed from the maternal genome and encode the maternal factors that are deposited into the developing oocytes in order to coordinate embryonic development before MBT [2]. We had previously explored the zebrafish egg transcriptome [3] and proteome [4] in order to gain further understanding of the maternal factors that contribute to good quality or developmentally competent eggs that result in high survival of progeny. However, large gaps still remain.

The forkhead box (Fox) proteins belong to a family of transcription factors that play important roles in cell growth, proliferation, survival, and cell death[5]. Many of these Fox proteins have been shown to be essential to the various processes of embryogenesis. In mammals, knockouts of several *fox* genes, including *foxa2*, *foxo1*, and *foxf1*, result in embryonic lethality due to defects in development of different organs ([5–7]). In reproduction, a recent transcriptomic study in the Nile tilapia, *Oreochromis niloticus*, showed that more than 50 *fox* genes were expressed in the gonads, and some of these, including *foxl2*, *foxo3*, and *foxr1*, were specific to XX females[8]. *foxl2* and its relatives are known to be key players in ovarian differentiation and oogenesis in vertebrates; it is essential for mammalian ovarian maintenance and through knockout experiments, it was demonstrated that *foxl2* is a critical regulator of sex determination by regulating ovary development and maintenance also in Nile tilapia, medaka, and zebrafish[9]. Further, *foxo3* was shown to be required for ovarian follicular development, and its knockout in mice led to sterility in female mutants due to progressive degeneration of the developing oocytes and lack of ovarian reserve of mature oocytes[10]. *foxr1* was also found to have sexually dimorphic expression in eels (*Anguilla anguilla* and *Monopterus albus*) and marine medaka (O*ryzias melastigma*) which was predominately observed in the ovaries[11–13]. However, despite these observational studies, the function of *foxr1* in vertebrates especially its role in reproduction remains unclear. Thus, in this study, we investigated the evolution of *foxr1* and its phylogenetic relationship in a wide range of vertebrate species, as well as its biological function using knockout zebrafish models created by the CRISPR/cas9 system in order to broaden our knowledge on the evolutionary origin of maternal-effect genes and the underlying mechanisms that contribute to reproductive success in vertebrates.

## Materials and Methods

### Protein databases

Since our model is based on the zebrafish, all gene/protein nomenclatures will be based on those of fish. The following amino acid data were extracted and investigated from the ENSEMBL database (http://www.ensembl.org/index.html): human, *Homo sapiens*; mouse, *Mus musculus*; rat, *Rattus norvegicus*; guinea pig, *Cavia porcellus*; pig, *Sus scrofa*; horse, *Equus caballus*; cow, *Bos taurus*; panda, *Ailuropoda melanoleuca*; opossum, *Monodelphis domestica*; Chinese softshell turtle, *Pelodiscus sinensis*; armadillo, *Dasypus novemcinctus*; frog, *Xenopus tropicalis*; fruit fly, *Drosophila melanogaster*; nematode, *Caenorhabditis elegans*; sea squirt, *Ciona intestinalis*; lamprey, *Petromyzon marinus*; coelacanth, *Latimeria chalumnae*; spotted gar, *Lepisosteus oculatus*; cod, *Gadus morhua*; fugu, *Takifugu rubripes*; medaka, *Oryzias latipes*; platyfish, *Xiphophorus maculatus*; stickleback, *Gasterosteus aculeatus*; tetraodon, *Tetraodon nigroviridis*; tilapia, *Oreochromis niloticus*; zebrafish, *Danio rerio*; and cave fish, *Astyanax mexicanus*. The bald eagle, *Haliaeetus leucocephalu*; penguin, *Pygoscelis adeliae*; crested ibis, *Nipponia nippon*; swan goose, *Anser cygnoides domesticus*; American alligator, *Alligator mississippiensis*; Chinese alligator, *Alligator sinensis*; python, *Python bivittatus*; central bearded dragon, *Pogona vitticeps*; frog, *Xenopus laevis*; medaka, *Oryzias latipes*; zebrafish, *Danio rerio*; northern pike, *Esox lucius*; rainbow trout, *Oncorhynchus mykiss*; coho salmon, *Oncorhynchus kisutch*; and Atlantic salmon, *Salmo salar*, protein sequences were extracted and investigated from the NCBI database (http://www.ncbi.nlm.nih.gov). Further, the following protein sequences were extracted from our previously established PhyloFish online database (http://phylofish.sigenae.org/index.html) [14] and analyzed along with the others: spotted gar, *Lepisosteus oculatus*; cod, *Gadus morhua*; bowfin, *Amia calva*; European eel, Anguilla anguilla; butterflyfish, *Pantodon buchholzi*; sweetfish, *Plecoglossus altivelis*; allis shad, *Alosa alosa*; arowana, *Osteoglossum bicirrhosum*; panga, *Pangasius hypophthalmus*; northern pike, *Esox lucius*; eastern mudminnow, *Umbra pygmae*; American whitefish, *Coregonus clupeaformis*; brook trout, *Salvelinus fontinalis;* rainbow trout, *Oncorhynchus mykiss*; European whitefish, *Coregonus lavaretus*; grayling, *Thymallus thymallus*; and European perch, *Perca fluviatilis*. These sequences are compiled in Supplemental Data 1.

### Phylogenetic analysis

The phylogenetic analysis was conducted using the Phylogeny.fr online program[15,16]. Amino acid sequences of 73 Foxr1, Foxr2, Foxn1, and Foxn3 proteins from the above-mentioned species were aligned using the MUSCLE pipeline, alignment refinement was performed with Gblocks, and then the phylogenetic tree was generated using the Maximum Likelihood method (PhyML pipeline) with 100 bootstrap replicates.

### Synteny analyses

Synteny maps of the conserved genomic regions of *foxr1* and *foxr2* were produced with spotted gar as the reference gene using PhyloView on the Genomicus v91.01 website (http://www.genomicus.biologie.ens.fr/genomicus-91.01/cgi-bin/search.pl).

### Quantitative real-time PCR (qPCR)

Tissue samples from 2 wildtype males and 3 wildtype females, and fertilized eggs at the one-cell stage from 32 wildtype couplings were harvested, total RNA was extracted using Tri-Reagent (Molecular Research Center, Cincinnati, OH) according to the manufacturer’s instructions. Reverse transcription (RT) was performed using 1 μg of RNA from each sample with the Maxima First Strand cDNA Synthesis kit (Thermo Scientific, Waltham, MA). Briefly, RNA was mixed with the kit reagents, and RT performed at 50°C for 45 min followed by a 5-min termination step at 85°C. Control reactions were run without reverse transcriptase and used as negative control in the qPCR study. qPCR experiments were performed with the Fast-SYBR GREEN fluorophore kit (Applied Biosystems, Foster City, CA) as per the manufacturer’s instructions using 200 nM of each primer in order to keep PCR efficiency between 90% and 100%, and an Applied Biosystems StepOne Plus instrument. RT products, including control reactions, were diluted 1/25, and 4 μl of each sample were used for each PCR. All qPCR experiments were performed in duplicate. The relative abundance of target cDNA was calculated from a standard curve of serially diluted pooled cDNA and normalized to *18S*, *β-actin*, and *EF1α*. transcripts. The primer sequences can be found in Supplemental Data 2. The tissue expression of *foxr1* was detected using the *foxr1* forward and reverse primers while the mutant form of *foxr1* in the CRISPR/cas9-mutated eggs was assessed with the mutant *foxr1* forward and reverse primers.

### RNA-seq

RNA-seq data were deposited into Sequence Read Archive (SRA) of NCBI under accession references SRP044781-84, SRP045138, SRP045098-103, and SRP045140-146. The construction of sequencing libraries, data capture and processing, sequence assembly, mapping, and interpretation of read counts were all performed as previously [14]. The number of mapped reads was then normalized for the *foxr1* gene across the 11 tissues using RPKM normalization.

### *In situ* hybridization (ISH)

Ovary samples were first fixed in 4% paraformaldehyde overnight, dehydrated by sequential methanol washes, paraffin-embedded, and sectioned to 7 µm thickness before being subjected to the protocol. The sections were deparaffinized and incubated with 10 µg/mL of proteinase K for 8 minutes at room temperature, followed by blocking with the hybridization buffer (50% formamide, 50 µg/mL heparin, 100 µg/mL yeast tRNA, 1% Tween 20, and 5X saline-sodium citrate [SSC]). The probe was diluted to 1 ng/µL in the hybridization buffer and incubated overnight at 55°C in a humidification chamber. The probes were synthesized by cloning a fragment of the *foxr1* gene into the pCRII vector using the cloning *foxr1* forward and reverse primers (Supplemental Data 2) and Topo TA Cloning kit (Invitrogen, Carlsbad, CA) as per the manufacturer’s protocol. The digoxigenin (DIG)-labeled sense and anti-sense probes were transcribed from Sp6 and T7 transcription sites, respectively, of the vector containing the cloned *foxr1* fragment and purified using 2.5M LiCl solution. The purity and integrity of the probes were verified using the Nanodrop spectrophotometer (Thermo Scientific) and the Agilent RNA 6000 Nano kit along with the Agilent 2100 bioanalyzer (Santa Clara, CA). The slides were then subjected to 2 washes each with 50% formamide/2X SSC, 2X SSC, and 0.2X SSC at 55°C followed by 2 washes with PBS at room temperature. The sections were subsequently blocked with blocking buffer (2% sheep serum, 3% bovine serum albumin, 0.2% Tween 20, and 0.2% Triton-X in PBS), and the anti-DIG antibody conjugated to alkaline phosphatase (Roche Diagnostics, Mannheim, Germany) was diluted by 1/500 and applied for 1.5 hours at room temperature. The sections were washed with PBS and visualized with NBT/BCIP (nitro blue tetrazolium/5-bromo-4-chloro-3-indolyl phosphate).

### CRISPR-cas9 genetic knockout

Fish used in this study were reared and handled in strict accordance with French and European policies and guidelines of the INRA LPGP Institutional Animal Care and Use Committee, which approved this study. CRISPR/cas9 guide RNA (gRNA) were designed using the ZiFiT[17,18] online software and were made against 2 targets within the gene to generate a genomic deletion of approximately 240 base pairs (bp) that spans the last exon which allowed the formation of a non-functional protein. Nucleotide sequences containing the gRNA were ordered, annealed together, and cloned into the DR274 plasmid. *In vitro* transcription of the gRNA from the T7 initiation site was performed using the Maxiscript T7 kit (Applied Biosystems) and of the cas9 mRNA using the mMESSAGE mMACHINE kit (Ambion/Thermo Scientific) from the Sp6 site, and their purity and integrity were assessed using the Agilent RNA 6000 Nano Assay kit and 2100 Bioanalyzer. Zebrafish embryos at the one-cell stage were micro-injected with approximately 30-40 pg of each CRISPR/cas9 guide along with purified cas9 mRNA. The embryos were allowed to grow to adulthood, and genotyped using fin clip and PCR that detected the deleted region. The full-length wildtype PCR band is 400 bp, and the mutant band with the CRISPR/cas9-generated deletion is approximately 160 bp. The PCR bands of the mutants were then sent for sequencing to verify the deletion. Once confirmed, the mutant females were mated with *vasa::gfp* males to produce F1 embryos, whose phenotypes were subsequently recorded. Images were captured with a Nikon AZ100 microscope and DS-Ri1 camera (Tokyo, Japan).

### Genotyping by PCR

Fin clips were harvested from animals under anesthesia (0.1% phenoxyethanol) and lysed with 5% chelex containing 100 µg of proteinase K at 55°C for 2 hrs and then 99°C for 10 minutes. The extracted DNA was subjected to PCR using Jumpstart Taq polymerase (Sigma-Aldrich, St. Louis, MO) and the *foxr1* forward and reverse primers that are listed in Supplemental Data 2.

### Statistical Analysis

Comparison of two groups was performed using the GraphPad Prism statistical software (La Jolla, CA), and either the Student’s t-test or Mann-Whitney U-test was conducted depending on the normality of the groups based on the Anderson-Darling test. A p-value < 0.05 was considered as significant.

## Results

### Phylogenetic analysis of Foxr1-related sequences

To date, there are six reported members of the *foxr*/*foxn* family (*foxn1*-*6*), of which *foxn5* and *foxn6* are also known as *foxr1* and *foxr2*, respectively. To examine the evolution of *foxr1*, we used a Blast search approach using the zebrafish Foxr1 protein sequence as query in various public databases to retrieve 73 protein sequences that could be related to this protein. All retrieved sequences are compiled in Supplemental Data 1. Of note, both Foxr1 and Foxr2 protein sequences were retrieved. In order to verify that the retrieved protein sequences (Supplemental Data 1) are homologous to zebrafish Foxr1, a phylogenetic analysis was performed. Based on the alignment of the retrieved vertebrate Foxr1-related sequences, and using Foxn1 and Foxn3 amino acid sequences as out-groups, a phylogenetic tree was generated (Fig 1). As shown in Fig 1, the common ancestor of the vertebrate Foxr1/Foxr2 diverged from Foxn1 and Foxn3, and these sequences were clearly observed as two separate clades belonging to actinopterygii (ray-finned fish) and sarcopterygii (lobe-finned fish and tetrapods). In addition, Foxr2 was found only in mammals with no homologs detected in actinopterygii as well as sauropsids and amphibians. Remarkably, despite the wide-ranging presence of Foxr1, no related sequences were observed in invertebrates and chondrichthyans (dogfish and sharks) as well as certain species such as chicken (*Gallus gallus*). On the other hand, several species showed two Foxr1 sequences including the salmonids, rainbow trout (*Oncorhynchus mykiss*) and brook trout (*Salvelinus fontinalis*), as well as northern pike (*Esox lucius*), cod (*Gadus morhua*), medaka (*Oryzias latipes*), and spotted gar (*Lepisosteus oculatus*). The presence of two related Foxr1 sequences in these species could be due to an independent gene duplication event that occurred in these species.

**Figure 1:**
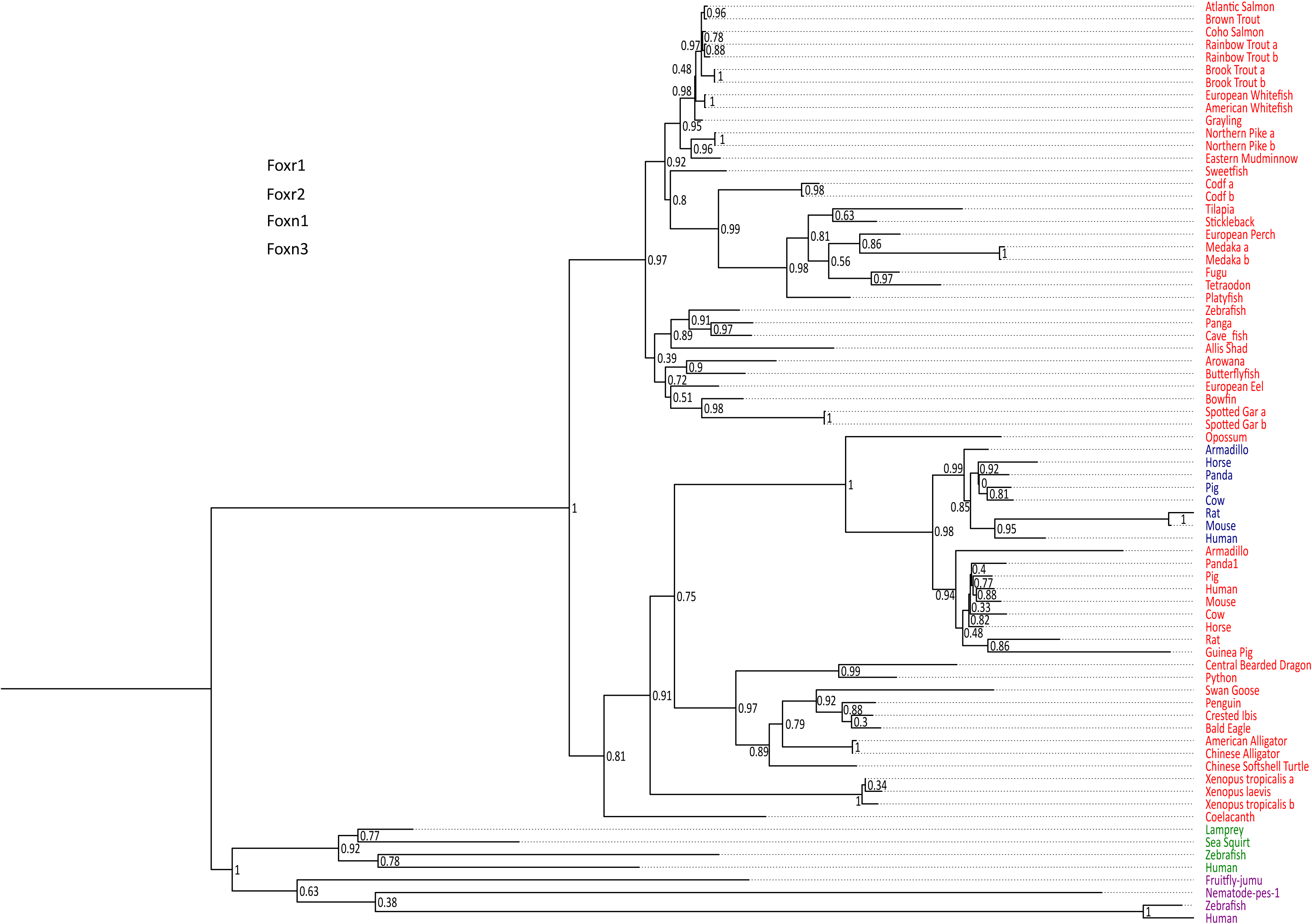
Phylogenetic tree of vertebrate Foxr1 and Foxr2 proteins. This phylogenetic tree was constructed based on the amino acid sequences of Foxr1 proteins (for the references of each sequence see Supplemental Data 1) using the Maximum Likelihood method with 100 bootstrap replicates. The number shown at each branch node indicates the bootstrap value (%). The tree was rooted using Foxn1 and Foxn3 sequences. The Foxr1 sequences are in red, Foxr2 sequences are in blue, those of Foxn1 are in green, and Foxn3 sequences are in purple.

Despite the previous report that stated that *foxr2* was absent in tilapia, stickleback, zebrafish, and medaka genomes, we retrieved Foxr2 protein sequences using the zebrafish Foxr1 peptide sequence as query. Thus, using zebrafish Foxr1 sequence as the reference protein, we subsequently compared its homology with the Foxr1 and Foxr2 sequences from mammals. As shown in Supplemental Data 3, there was 29-37% positivity and 41-53% similarity between all sequences, and there did not appear to be any difference in homology between zebrafish Foxr1 and mammalian Foxr1 and Foxr2 sequences. Further, there was 47-60% positivity and 59-77% similarity between mammalian Foxr1 and Foxr2 sequences, indicating that these two proteins are highly similar and probably diverged quite late in evolution.

### Synteny analysis of *foxr1* and *foxr2* genes in vertebrates

In order to further understand the origin of the *foxr1* and *foxr2* genes in vertebrates, we performed a synteny analysis of their neighboring genes in representative vertebrate genomes using the basal actinopterygian, spotted gar, as the reference genome and the Genomicus online database (Fig 2). We found that between the spotted gar and mammals, there was conserved synteny of the *foxr1*, *upk2*, *ccdc84*, *rps25*, *trappc4*, *slc37a4*, and *ccdc153* loci in their genomes. In the frog (*Xenopus tropicalis*) genome, the *foxr1*, *ccdc153*, *cbl*, *mcam*, and *c1qtnf5* loci were conserved, while in Coelacanth, *foxr1*, *ccdc84*, *rps25*, *trappc4*, *slc37a4*, *cbl*, *ccdc153*, *mcam*, *c1qtnf5*, as well as *rnf26* loci were found in the same genomic region as those of the spotted gar. However, amongst the actinopterygians, there was lower conservation of synteny; in zebrafish and cave fish, the *foxr1*, *ccdc84*, and *mcam* loci were conserved while in the other ray-finned fish species, only the *foxr1* loci was found. We further analyzed the *foxr2* sequences that were found only in mammals, and we demonstrate here that they were all observed on the X chromosome with no apparent conserved synteny of neighboring genes to those found in the spotted gar. Our overall analyses suggest that all the *foxr*-related sequences that were found were homologs, and the *foxr* gene in fish species probably derived from the ancestral *foxr1* gene. Although there was the same degree of protein homology between zebrafish Foxr1 and mammalian Foxr1 and Foxr2 sequences, the phylogenetic tree and synteny analyses showed a clear distinction between them, and the *foxr2* gene probably derived from a later single gene duplication or transposon event as previously suggested[19].

**Figure 2:**
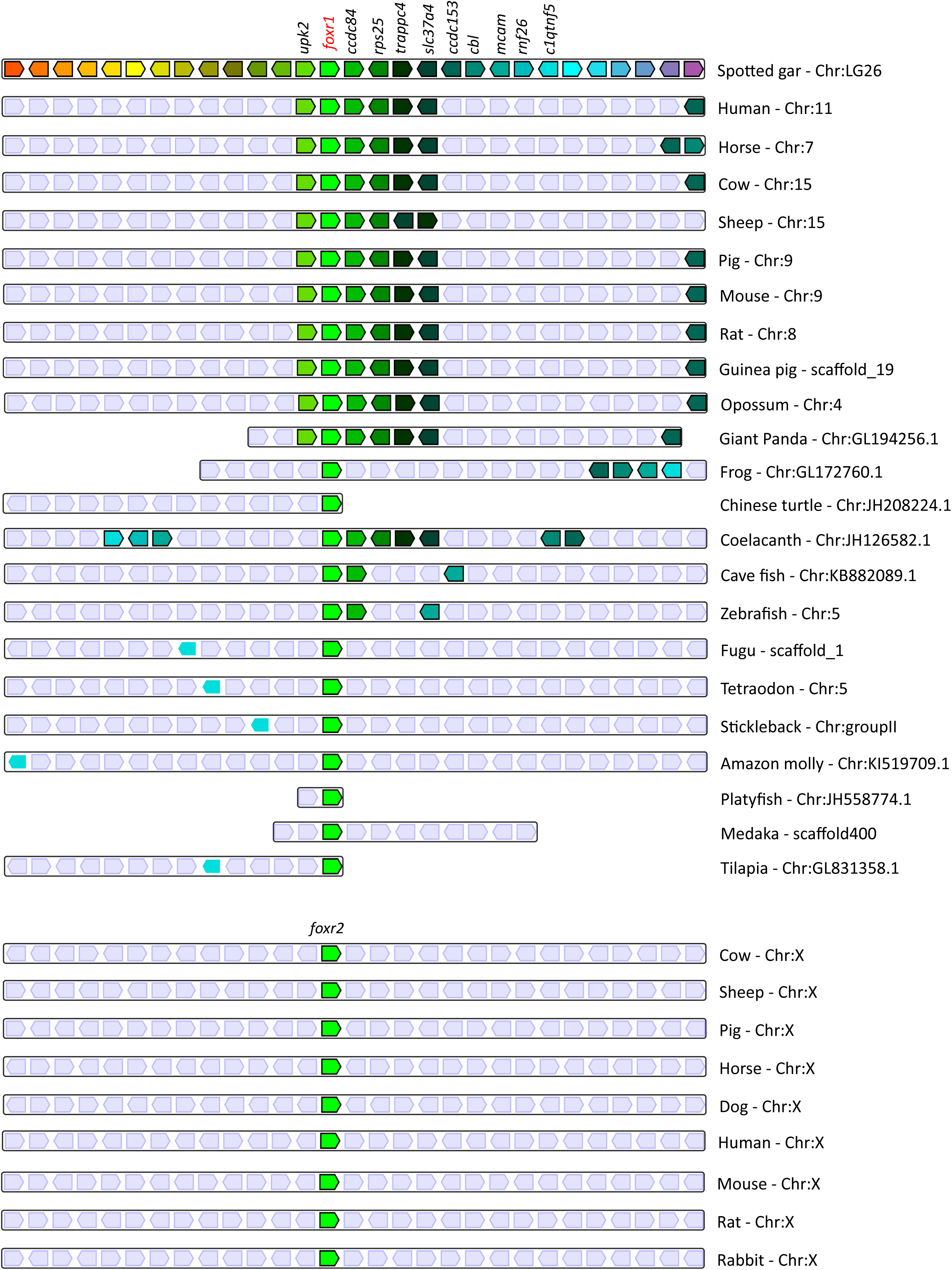
Conserved genomic synteny of *foxr1* genes. Genomic synteny maps comparing the orthologs of *foxr1, foxr2*, and their neighboring genes, which were named after their human orthologs according to the Human Genome Naming Consortium (HGNC). Orthologs of each gene are shown in the same color, and the chromosomal location is shown next to the species name. *foxr1* orthologs are boxed in red while *foxr2* orthologs are boxed in blue.

### Expression profiles of *foxr1*

We next focused our efforts on *foxr1 s*ince it has previously been shown in eel, tilapia, and medaka to be gonad specific and thus may have specific functions in reproduction. In order to investigate the potential functions of *foxr1*, we explored its tissue distribution using two different approaches, qPCR and RNA-seq, the latter of which was obtained from the PhyloFish online database [14]. In zebrafish, we observed from both sets of data that *foxr1* mRNA was predominantly expressed in the ovary and unfertilized egg (Fig 3A and 3B). By ISH, we also demonstrated that *foxr1* transcripts were highly expressed in the ovary in practically all stages of oogenesis (Fig 3C-E; negative controls, Fig 3F-H).

**Figure 3:**
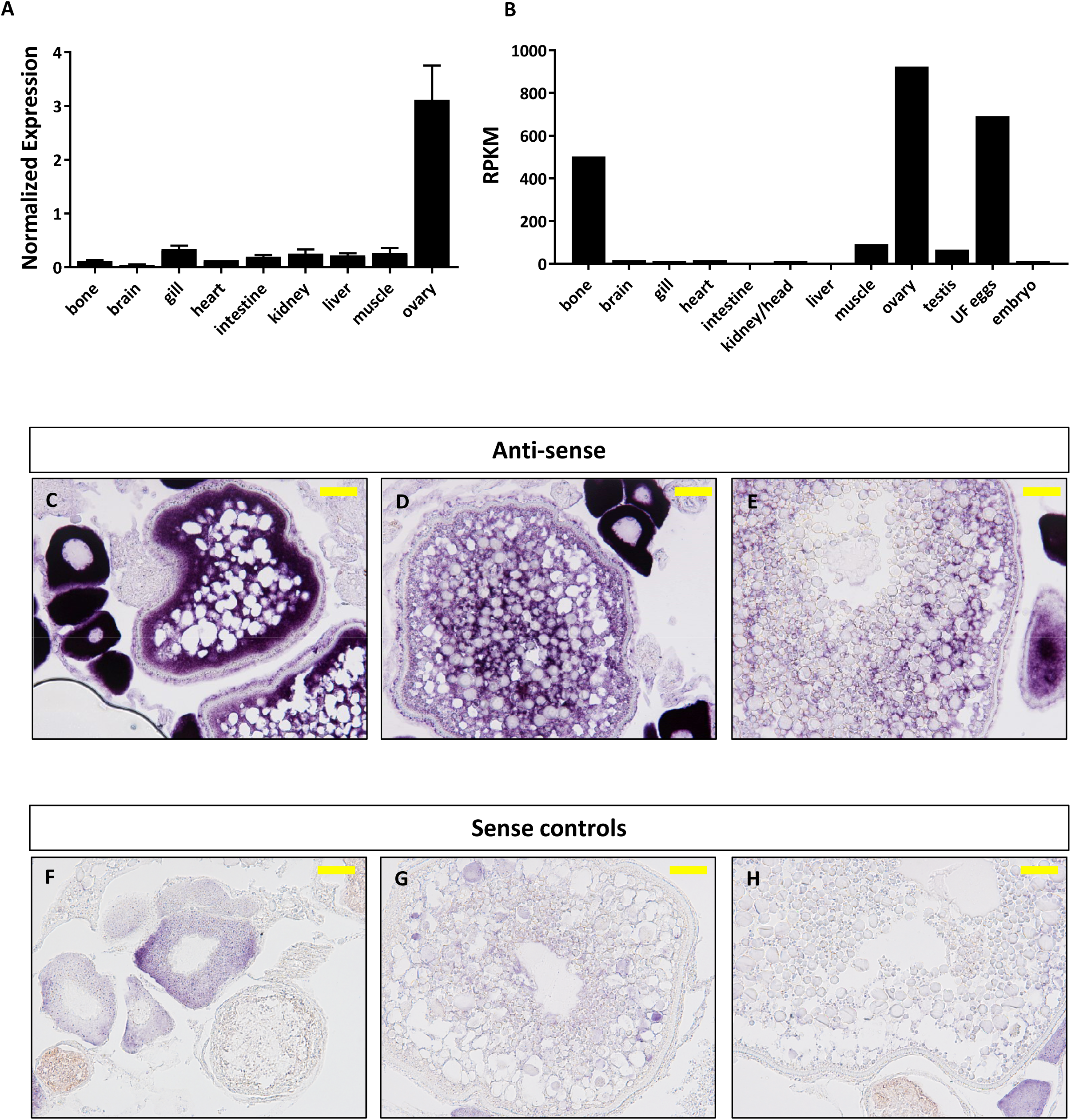
Expression profile of *foxr1* in zebrafish. Tissue expression analysis of *foxr1* mRNA in zebrafish (**A)** by quantitative real-time PCR (qPCR) and **(B)** RNA-seq. Expression level by qPCR is expressed as a normalized value following normalization using *18S*, *β-actin*, and *EF1α*. expression while that by RNA-seq is expressed in read per kilobase per million reads (RPKM). Tissues were harvested from 3 to 4 wildtype zebrafish individuals. **(C-H)** *In situ* hybridization was performed for *foxr1* in zebrafish ovaries from wildtype females. Positive staining is demonstrated using the anti-sense probe against *foxr1* (Fig 3C-E) in blue with 5-bromo-4-chloro-3-indolyl-phosphate/nitro blue tetrazolium as substrate. The negative control was performed with the sense probe (Fig 3F-H). 20X magnification; bars denote 90 µm. N=5 each for *foxr1* mutant and control.

### Functional analysis of *foxr1* in zebrafish

To understand the role of *foxr1* during oogenesis and early development, we performed functional analysis by genetic knockout using the CRISPR/cas9 system. One-cell staged embryos were injected with the CRISPR/cas9 guides that targeted *foxr1* and allowed to grow to adulthood. Mosaic founder mutant females (F0) were identified by fin clip genotyping and subsequently mated with *vasa::gfp* males, and embryonic development of the F1 fertilized eggs was recorded. Since the mutagenesis efficiency of the CRISPR/cas9 system was very high, as previously described [20,21], the *foxr1* gene was sufficiently knocked-out even in the mutant mosaic F0 females. This was evidenced by the substantially lower transcript level of *foxr1* in the F1 embryos as compared to those from control pairings (Fig 4A). Thus, the phenotypes of *foxr1* (n=5) mutants could be observed even in the F0 generation. Since none of the mutated genes were transmissible to future generations neither through the male nor the female (ie. all the surviving embryos were WT), therefore, all of our observations were obtained from the F0 generation.

**Figure 4:**
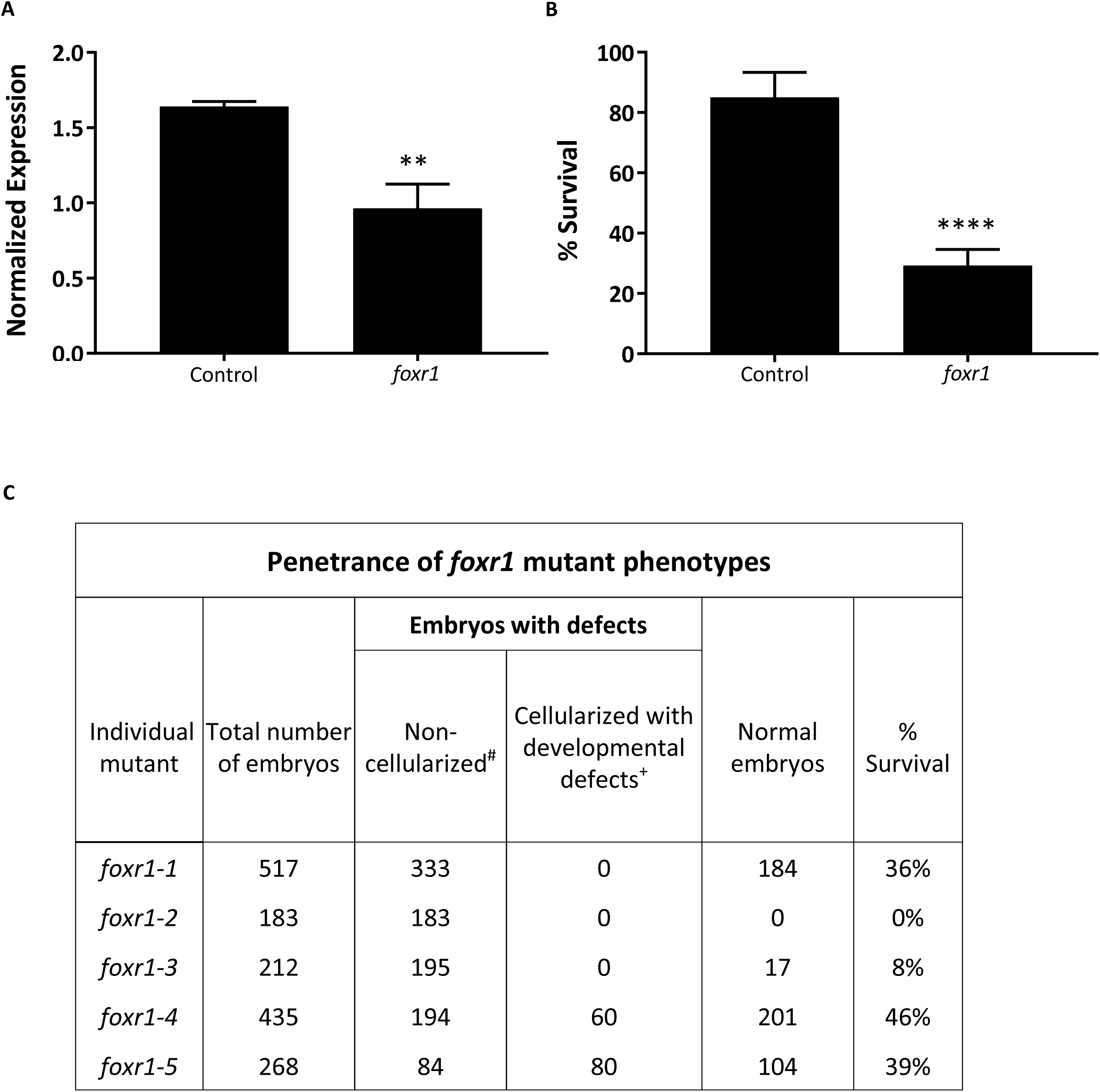
CRISPR/cas9 knockout of *foxr1* in zebrafish. **(A)** Normalized expression level of *foxr1* transcript by quantitative real-time PCR (qPCR) in the fertilized zebrafish eggs from crosses between *foxr1* mutant females and *vasa::gfp* males. **(B)** Developmental success (% survival) at 24 hours post-fertilization (hpf) as measured by the proportion of fertilized eggs that underwent normal cell division and reached normal developmental milestones based on Kimmel et al. [30] from crosses between *foxr1* mutant females and *vasa::gfp* males. **(C)** Penetrance of *foxr1* mutant phenotypes in the F1 eggs between crosses of *foxr1* mutant females and *vasa::gfp* males. The graph demonstrates representative data from a single clutch from each mutant female. ^#^Embryos did not develop at all (please refer to Fig 5E-H). ^+^Embryos had a partially cellularized blastodisc that was sitting atop an enlarged syncytium (please refer to Fig 7I-L).qPCR data were normalized to *18S*, *β-actin*, and *EF1α*.. N=5 each for *foxr1* mutant and control. All assessments were performed from at least 3 clutches from each mutant. ** p<0.01, ****p<<0.0001 by Mann-Whitney U-test. Control = eggs from crosses of wildtype females with *vasa::gfp* males; *foxr1* = eggs from crosses of *foxr1* mutant females with *vasa::gfp* males.

We observed that most of the embryos from the *foxr1* mutant females had a very low developmental success at 24 hpf (25.2±5.5% vs. 85.1±8.3% in controls; p<0.0001) (Fig 4B). The penetrance of the mutation in the mutant females is demonstrated in Fig 4C, and it was observed that 3 of the mutants produced abundant non-developing eggs that remained non-cellularized, reflecting their failure to undergo cell division (Fig 5E-H). The eggs derived from these 3 *foxr1* mutant females did not undergo any cell division at 1 hpf and continued to display a complete lack of development up to 8 hpf. By 24 hpf, these non-developing eggs that failed to divide were all dead. In addition, two of the mutants produced developmentally incompetent eggs with two phenotypes; those with a non-cellularized morphology (Fig 5E-H), and another population that developed albeit with an abnormal morphology (Fig 5I-L). These fertilized and developing embryos were structurally abnormal, with unsmooth and irregularly-shaped yolk as well as asymmetrical cell division that culminated into a blastodisc with a group of cells on top of an enlarged syncytium (arrow). These eggs underwent developmental arrest at around 4 hpf or the MBT and appeared to regress with further expansion of the syncytium (Fig 5J-K) until death by 24 hpf. This phenotype was also observed previously by us in *npm2b* mutant-derived eggs[22].

**Figure 5:**
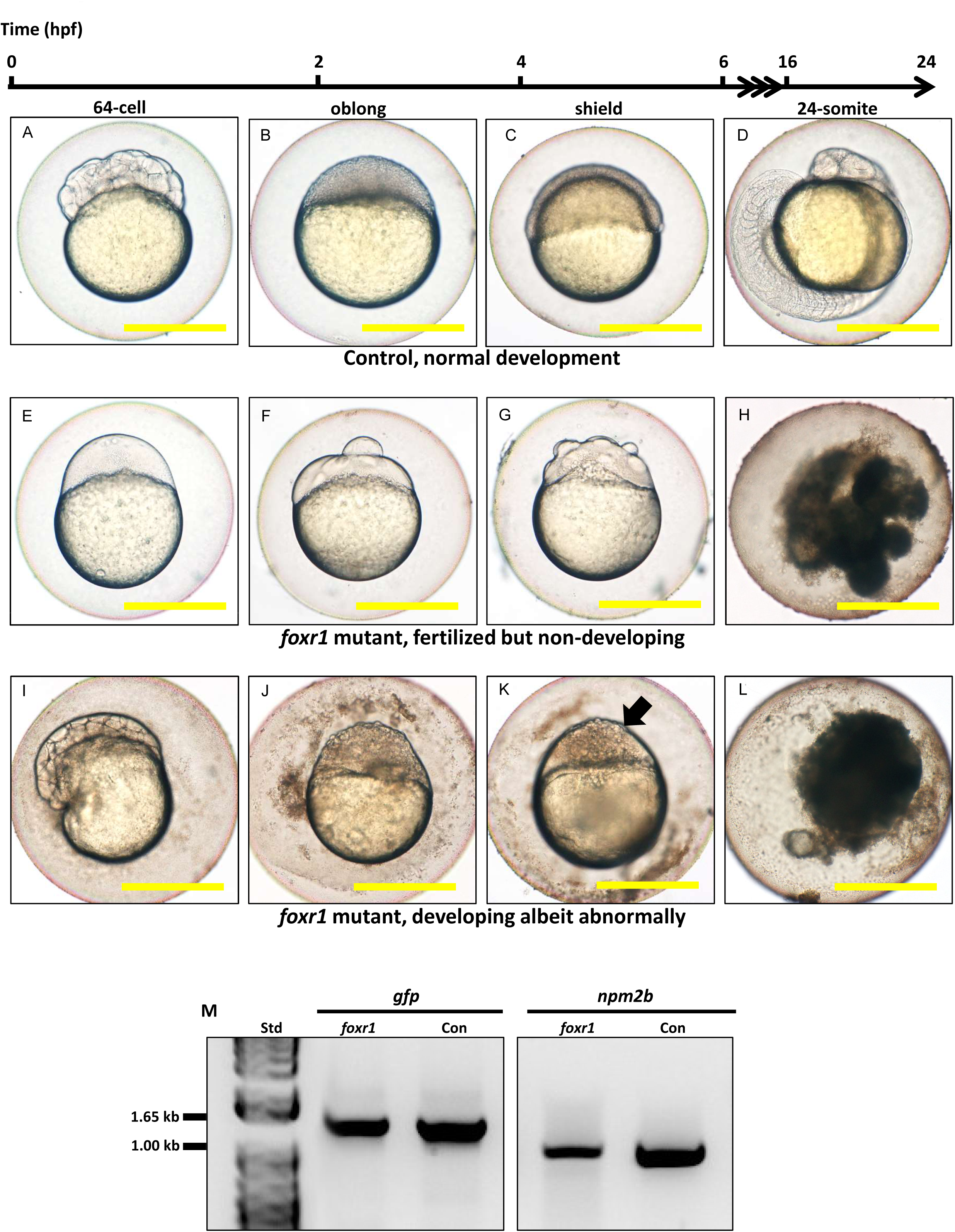
Effect of *foxr1* deficiency on zebrafish embryogenesis. Representative images demonstrating development of fertilized eggs from crosses between control **(A-D)** and *foxr1* **(E-L)** females and *vasa::gfp* males from 2-24 hours post-fertilization (hpf). In the control eggs, the embryos were at 64-cell **(A)**, oblong **(B)**, germ ring **(C)**, and 24-somite **(D)** stages according to Kimmel et al [30]. Eggs from *foxr1* mutant females were non-developing with a non-cellularized morphology **(E-H)** or developing with an abnormal morphology **(I-L)**. **(A, E, I)** = images taken at 2 hpf; **(B, F, J)** = images taken at 4 hpf; **(C, G, K)** = images taken at 6 hpf; **(D, H, L)** = images taken at 24 hpf. Scale bars denote 500 µm. The arrow demonstrates an abnormally cellularized blastodisc that was sitting atop an enlarged syncytium. **(M)** Genotypic analysis of the eggs from crosses of *foxr1* mutant females and *vasa::gfp* males to determine fertilization status. The *gfp* and *vasa* primers produced a band that was 1333 base pairs in size. Detection of the *npm2b* gene (band size = 850 base pairs) was used as a control. Con = eggs from crosses of wildtype females with *vasa::gfp* males; *foxr1* = eggs from crosses of *foxr1* mutant females with *vasa::gfp* males. N=5 each for *foxr1* mutant and control.

The observed phenotype of the *foxr1* mutant-derived uncellularized eggs was very similar to previously described unfertilized eggs [23]. Thus, the foxR1 mutant females were mated with *vasa::gfp* males, and the genotype of their progeny was assessed for the presence of the *gfp* gene, which would only be transmitted from the father since the mutant females did not carry this gene. We found that these uncellularized eggs from the *foxr1* mutant females did indeed carry the *gfp* gene (Fig 4M) which indicated that they were fertilized, but were arrested from the earliest stage of development and did not undergo any cell division. These novel findings showed for the first time that *foxr1* is essential for the developmental competence of zebrafish eggs, and is therefore a crucial maternal-effect gene.

In order to delve into the possible mechanisms that may be involved in the reduced reproductive success of the *foxr1* mutants, we investigated the expression levels of *p21*, *p27*, and *rictor*, which were previously reported to be repressed by the Foxr1 transcription factor in mice (Santo et al, 2012). We found that there was substantially increased expression of *p21* (4.83±1.09 vs 0.25±0.03 in controls; p<0.0022) while that of *rictor* was significantly decreased (0.83±0.11 vs 1.81±0.23 in controls; p<0.0007) in the *foxr1* mutant-derived eggs as compared to eggs produced by wildtype females (Fig 6A-C). These results were in line with a growth arrested phenotype that was observed in the uncellularized and developmentally challenged eggs from the *foxr1* mutant females.

**Figure 6:**
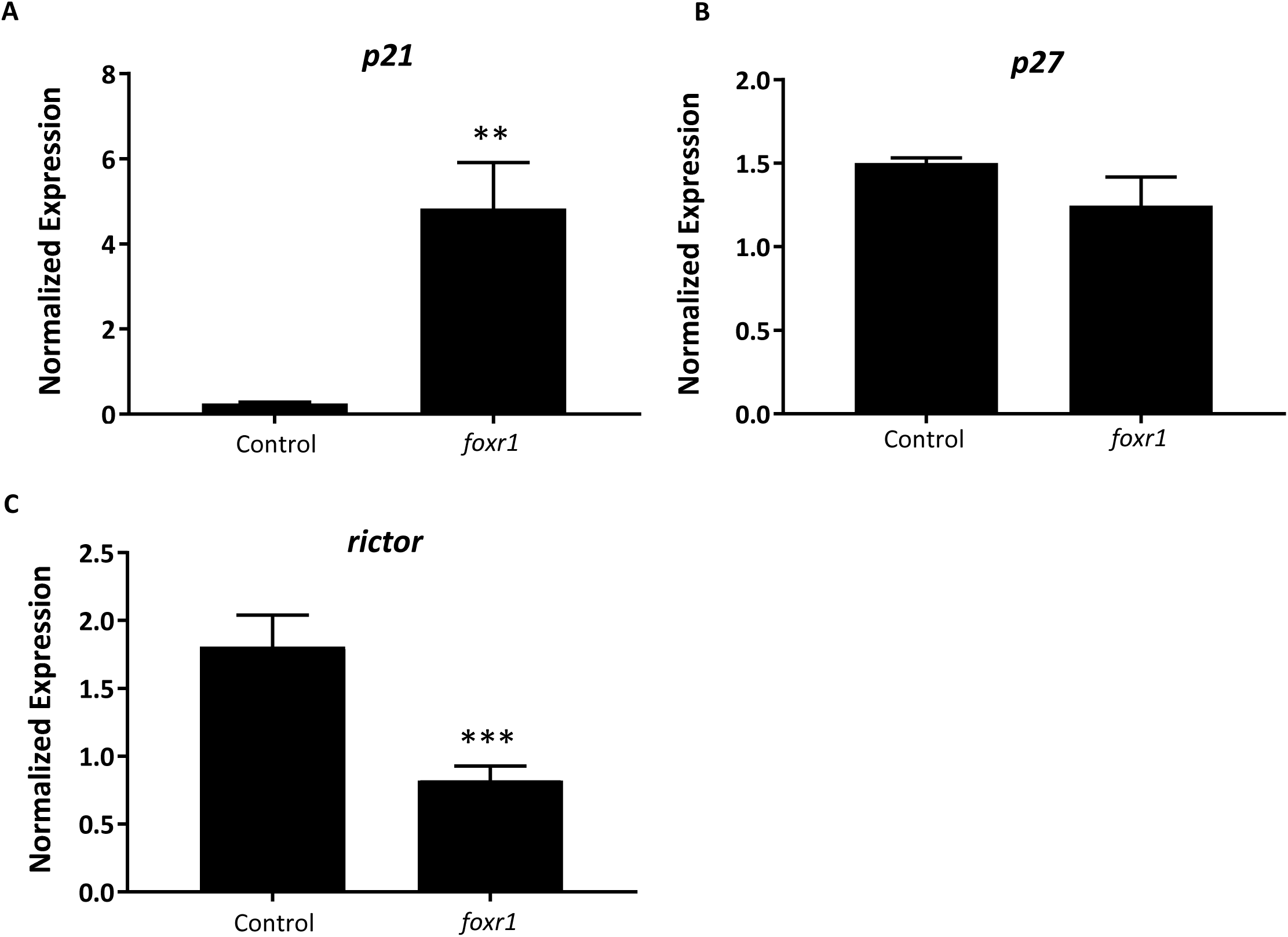
Expression profiles of *p21*, *p27*, and *rictor* in eggs from *foxr1* mutant females. Fertilized eggs from *foxr1* mutant females were subjected to qPCR for examination of the transcript levels of *p21*, *p27*, and *rictor*. Data were normalized to *18S*, *β-actin*, and *EF1α*.. N=5 each for *foxr1* mutant and control, at least two clutches were used from each animal, and each experiment was performed in duplicate. **p<0.01, ***p<0.001 by Mann-Whitney U-test.

## Discussion

In this study, we first investigated the evolutionary history of *foxr1* in order to gain perspective into its phylogenetic relationship among homologs from a wide range of species and to clarify its origins. Using the zebrafish protein sequence as query to search for homologs in other species, we retrieved Foxr1 sequences from a broad variety of vertebrates, including actinopterygii, sarcopterygii, and sauropsids which suggested the essentialness of this protein in most vertebrates. We also retrieved Foxr2 sequences due to its high similarity to the zebrafish Foxr1 peptide (Supplemental Data 3), although we and others demonstrated that the *foxr2* gene is absent from all actinopterygii and sauropsid species, and can only be found in mammals. Evidence from the phylogenetic analyses showed a clear distinction in derivation of the actinopterygian *foxr1* and the mammalian *foxr2;* the divergence of the ancestral *foxr1* gene in actinopterygii from that of the sarcopterygii and sauropsids occurred quite early in evolution, while the divergence of mammalian *foxr1* and *foxr2* is a much more recent event (Fig1). Further, the synteny analysis (Fig2) showed that there was much conservation of genomic synteny surrounding the *foxr1* loci between the basal actinopterygian, spotted gar, and actinopterygii and sauropsids, while the neighboring loci around the *foxr2* were completely different in comparison to those next to *foxr1* which suggested that *foxr2* originated from a recent gene duplication or transposon event as previously proposed[19]. We also found that in a small subset of species [rainbow trout (*Oncorhynchus mykiss*) and brook trout (*Salvelinus fontinalis*), as well as northern pike (*Esox lucius*), cod (*Gadus morhua*), medaka (*Oryzias latipes*), and spotted gar (*Lepisosteus oculatus*)], two Foxr1 sequences were observed. This suggested that a single gene duplication event may have occurred in these species and subsequent gene losses after the multiple genome duplication events such as the teleost-specific whole genome duplication (TGD) and salmonid-specific whole genome duplication (SaGD) occurred in the teleosts. It is also possible that *foxr1* was duplicated in the ancestral actinopterygii and subsequent gene losses in bowfin as well as in the teleosts especially following the multiple gene duplication events such as the teleost-specific whole genome duplication (TGD) and salmonid-specific whole genome duplication (SSGD). Thus, it appeared that TGD and SSGD did not impact the current *foxr1* gene diversity because in most species, only one *foxr1* gene was retained. The presence of two *foxr1* genes in the above-mentioned species could also be due to independent and phylum-specific gene retention or independent gene duplication events that occurred only in these species. Further analyses on the two copies of *foxr1* in these species are warranted in order to verify the functionality of both genes.

The essentialness of *foxr1* was suggested by the wide-ranging presence of this gene in most vertebrates and the retention of a single copy in most teleosts despite multiple whole genome duplication events, but its biological function is still largely unknown. Previous reports have demonstrated the predominant expression of *foxr1* mRNA in the ovary of medaka, eel, and tilapia[8,11,13], but in the male germ cells and spermatids in mouse and human[24]. It was further shown to be abundantly expressed in the early cleavage and gastrula stages of Xenopus embryos, but absent in post-gastrula stages due to rapid degradation of its mRNA, indicating that it is a maternally-inherited transcript[25]. Thus, the *foxr1* gene may play different roles in reproduction in teleost fish/amphibians and mammals, suggesting that *foxr2* in mammals may have evolved to have comparable functions to the teleost/amphibian *foxr1*. Future studies to test this are necessary to confirm the function of *foxr2.* To confirm these results found in other teleosts in zebrafish, we first examined the expression profile of *foxr1* in various tissues, and we showed by qPCR as well as by RNA-seq that there was also an ovarian-specific expression of *foxr1* and negligible amount in the testis as in the other fish species. By ISH, we found that the *foxr1* transcript was progressively stored in the growing oocytes from the very early stages (Fig 3C-D, arrows) to later staged oocytes (Fig 3D-E), and could be found abundantly in mature fertilized eggs (Fig 3B and Fig 4A). These results demonstrated that *foxr1* is one of the maternal products that is deposited into the developing oocytes during oogenesis in zebrafish.

Having established that *foxr1* was indeed a maternal factor, we investigated its function via mutagenic analysis with CRISPR/cas9. We used the F0 mosaic mutant females that were shown to have a decreased level of *foxr1* mRNA for analysis due to the difficulty in transmitting the mutated *foxr1* gene to future generations as both the F0 *foxr1* mutant females and males produced mostly non-viable progeny, and the surviving descendents were all of wildtype genotype. This may be due to the efficiency of the CRISPR/cas9 mutagenic system in knocking out the *foxr1* gene very early on during the development of the animal. We found that the *foxr1* mutant females produced bad quality eggs, and the developmental success of their progeny was very low, similar to that of *foxl2* and *foxo3* mutants. Thus, it is likely that *foxr1* is also required for proper ovarian development and function. Further, we found that the *foxr1* mutant-derived eggs were non-cellularized and did not undergo subsequent cell division despite being fertilized. This suggested that their defect did not lie in the capability to be fertilized, as seen in *slc29a1a* and *otulina* mutants [3], but in the cell cycle and proliferation processes. Thus, we investigated the expression profiles of *p21*, *p27*, and *rictor*, which are all cell cycle and cell survival regulators, since Santo et al had previously knocked down *foxr1* using short hairpin RNAs in mammalian cells and found it to be a transcriptional repressor of them[26]. In this report, we also observed a dramatic increase in *p21* transcript in the eggs from *foxr1* mutant females, although the expression of *p27* was unchanged, while that of *rictor* was decreased. Both *p21* and *p27* are well known cell cycle inhibitors, and *rictor* is a component of the mTOR (mammalian target of rapamycin) complex that is a major regulator of cell growth and proliferation[27,28]. In fact, mitogens or some survival signal activates a survival cascade, such as the PI3K/Akt pathway, which is activated by the rictor-mTOR complex and promotes cell growth through repression of the negative cell cycle modulators, including *p21* and *p27*[29]. Thus, our findings were in line with a phenotype of growth arrest and anti-proliferative effects as seen in our eggs derived from *foxr1* mutant females. The different results that we observed as compared to those from Santo et al were probably due to species- and cell type-specific effects.

In this study, we showed that *foxr1* are found in a wide-range of vertebrates and are homologous to the *foxr1* genes found in other species. In teleosts, *foxr1* expression is found predominately in the ovary while in mammals, it appears to be specific to the male germline. We also found that *foxr1* is a novel maternal-effect gene and is highly expressed in the developing oocytes as well as accumulated in mature eggs to be used in early embryogenesis. Maternally-inherited *foxr1* is required for the first few cleavages after fertilization for proper cell growth and proliferation via *p21* and *rictor*, since deficiency in *foxr1* leads to either complete lack of or abnormal cell division culminating to early death in the fertilized egg. Thus, the results of this study establishes a link between egg quality and the control of early cell cycle and the mTOR patway via the potential transcriptional factor, foxR1.

## Author contributions

CTC performed the experiments and analyses, and drafted the manuscript; AP provided technical assistance in fish husbandry; YG participated in data analysis and manuscript preparation; and JB conceived and supervised the study and participated in data analysis and manuscript preparation.

## Acknowledgements

This work was supported by an ANR grant, Maternal Legacy (ANR-13-BSV7-0015), to JB. The authors would like to thank all the members of the LPGP laboratory for their assistance.

## Supporting Information

Supplemental Data 1. Foxr1-related protein sequences used in the phylogenetic analysis.

Supplemental Data 2. qPCR and PCR primer sequences.

Supplemental Data 3. Percentage homology between Foxr1 and Foxr2 protein sequences.

